# An evolutionarily ancient Fatty Acid Desaturase is required for the synthesis of hexadecatrienoic acid, which is the main source of the bioactive jasmonate in *Marchantia polymorpha*

**DOI:** 10.1101/2021.09.06.459162

**Authors:** Gonzalo Soriano, Sophie Kneeshaw, Guillermo Jimenez-Alemán, Angel M. Zamarreño, José Manuel Franco-Zorrilla, Mª Fernanda Rey-Stolle Valcarce, Coral Barbas, Jose M. García-Mina, Roberto Solano

## Abstract

Jasmonates are fatty acid derived hormones that regulate multiple aspects of plant development, growth and stress responses. Bioactive jasmonates, defined as the ligands of the conserved COI1 receptor, differ between vascular plants and bryophytes (using jasmonoyl-L-isoleucine; JA-Ile and dinor-12-oxo-10,15(*Z*)-phytodienoic acid; dn-OPDA, respectively). Whilst the biosynthetic pathways of JA-Ile in the model vascular plant *Arabidopsis thaliana* have been elucidated, the details of dn-OPDA biosynthesis in bryophytes are still unclear. Here, we identify an ortholog of *Arabidopsis* Fatty Acid Desaturase 5 (AtFAD5) in the model liverwort *Marchantia polymorpha* and show that FAD5 function is ancient and conserved between species separated by more than 450 million years of independent evolution. Similar to AtFAD5, MpFAD5 is required for the synthesis of 7*Z*-hexadecenoic acid. Consequently, in Mp*fad5* mutants the hexadecanoid pathway is blocked, dn-OPDA levels almost completely depleted and normal chloroplast development is impaired. Our results demonstrate that the main source of dn-OPDA in *Marchantia* is the hexadecanoid pathway and the contribution of the octadecanoid pathway, i.e. from OPDA, is minimal. Remarkably, despite extremely low levels of dn-OPDA, MpCOI1-mediated responses to wounding and insect feeding can still be activated in Mp*fad5*, suggesting that dn-OPDA is not the only bioactive jasmonate and COI1 ligand in *Marchantia*.

## INTRODUCTION

Jasmonates are fatty acid (FA) derived phytohormones that play a critical role in plant development and survival. They regulate responses to a wide range of stresses, both biotic (e.g. herbivory and necrotrophic pathogen infection) and abiotic (e.g. mechanical wounding, drought, or heat); (Wasternack & Feussner 2018, Howe *et al*., 2018; Monte *et al*., 2020). Moreover, they are involved in a multitude of developmental processes such as senescence, growth, flower development and fertility (Wasternack & Feussner 2018, Howe *et al*., 2018).

Most jasmonate-responsive processes are mediated by the COI1-JAZ co-receptor complex, where COI1 (Coronatine Insensitive 1) is the F-box component of an E3-type ubiquitin ligase, and JAZ (Jasmonate-ZIM domain) proteins are repressors of downstream transcription factors (TFs) (Chini *et al*., 2007; Thines *et al*., 2007; Sheard *et al*., 2010; Howe *et al*., 2018). During the activation of jasmonate signalling, a bioactive jasmonate triggers the interaction of COI1 and JAZ by acting as a “molecular glue” between both proteins, thus forming the co-receptor complex (Katsir *et al*., 2008; Fonseca *et al*., 2009; Sheard *et al*, 2010). Through its interaction with COI1, the JAZ repressor is ubiquitinated and then degraded by the proteasome, resulting in a de-repression of jasmonate-related TFs, including the jasmonate-responsive master regulator, MYC2 (Chini *et al*., 2007, Lorenzo *et al*, 2004, Fernandez-Calvo *et al*., 2011; Qi *et al*. 2015; Chini *et al*., 2016; Howe *et al*., 2018). In addition to this, through the electrophilic properties of their cyclopentanone rings, jasmonates also regulate a subset of genes through a COI1-independent pathway. It has recently been shown that this pathway activates heat stress (HS) responses and contributes to basal thermotolerance (Farmer & Mueller, 2013; Monte *et al*., 2020).

The octadecanoid pathway has been described as the major source of jasmonates biosynthesis in plants (Sup. Fig. **S1**). α-Linolenic acid (ALA, 18:3n3) is released from chloroplast membranes, and subjected to the consecutive actions of lipoxygenase, allene oxide synthase (AOS) and allene oxide cyclase (AOC) enzymes, to produce 12-oxo-phytodienoic acid (ODPA). This compound is transported to the peroxisome where it is converted into jasmonic acid (JA) via OPDA reductase 3 (OPR3)-mediated reduction and three successive rounds of β-oxidation. In some plant species, such as *Arabidopsis thaliana*, a parallel JA biosynthesis route exists, the hexadecanoid pathway, in which hexadecatrienoic acid (HTA, 16:3n3) undergoes the same processing as ALA (18:3n3) to produce a molecule structurally similar to OPDA, dinor-12-oxo-phytodienoic acid (dn-OPDA; Sup. Fig. **S1**; Weber *et al*., 1997). Furthermore, in *Arabidopsis*, analysis of *opr3* mutants elucidated an OPR3-independent pathway in which OPDA suffers three β-oxidations to produce dn-OPDA, tetranor-OPDA (tn-OPDA) and then 4,5-ddh-JA, which is finally reduced to JA by cytoplasmic OPRs (Chini *et al*., 2018).

In vascular plants, JA is conjugated to L-isoleucine (Ile) by JAR1 (JASMONATE RESISTANT 1), giving rise to the bioactive form of the hormone, (+)-7-*iso*-JA-Ile (JA-Ile; Staswick and Tiryaki, 2004; Thines *et al*., 2007; Fonseca *et al*., 2009; Sheard *et al*., 2010). Bryophytes, however, do not synthesise JA-Ile and dn-OPDA acts as the bioactive jasmonate. dn-OPDA binds a conserved COI1-JAZ co-receptor and regulates similar responses to those activated by JA-Ile in vascular plants (Monte *et al*., 2018; 2019; 2020; Peñuelas *et al*., 2019).

In plants, the synthesis of polyunsaturated fatty acids (PUFAs) such as ALA (18:3n3) and HTA (16:3n3) is carried out by a protein family known as fatty acid desaturases (FADs), which introduce a double bond regiospecifically in the FA backbone. FADs show high substrate specificity and are active in different subcellular compartments such as the endoplasmic reticulum (ER; the eukaryotic pathway) or in the chloroplasts (the prokaryotic pathway) (Fatiha, 2019; He *et al*., 2020).

In contrast to ALA (18:3n3), which can be synthesized in both the ER (by FAD3) (Arondel *et al*., 1992) and the plastids (by FAD7 and FAD8), HTA (16:3n3) is exclusively synthesized in the chloroplast and its production requires a specific FAD, the FAD5 (Weber *et al*., 1997). In *Arabidopsis*, this enzyme has been described as a plastidial Δ^7^-desaturase undertaking the first desaturation of palmitic acid to render 7*Z*-hexadecenoic acid (HD; a Δ^7^-16:1 FA that in this particular case matches a 16:1n9) (Heilmann *et al*., 2004a). After this first dehydrogenation, FAD6 and FAD7/FAD8 undertake two consecutive desaturation reactions to create Δ^7,10^-hexadecadienoic (HAD, 16:2n6) and HTA (16:3n3), respectively (Browse *et al*., 1989; Gibson *et al*., 1994; McConn *et al*., 1994).

Although the biosynthesis of PUFAs and dn-OPDA has been well studied in vascular plants (Weber *et al*., 1997; Chini *et al*., 2018; Fatiha, 2019; He *et al*., 2020), details of their production in bryophytes remain to be elucidated. It has been suggested that in bryophytes, as in *Arabidopsis*, dn-OPDA can be synthesized from both octadecanoid and hexadecanoid pathways. Indeed, in the model liverwort *Marchantia polymorpha* the addition of labelled ALA (18:3n3) or HTA (16:3n3) results in the accumulation of labelled dn-OPDA (Monte *et al*., 2018). However, it is unclear if the conversion of ALA/HTA into dn-OPDA occurs *in vivo* in untreated plants, and the enzymes involved in this process have not yet been characterised.

Here, we have elucidated the biosynthetic pathway of dn-OPDA in the model bryophyte *Marchantia*. We identified the *Marchantia* ortholog of AtFAD5 and generated a loss-of-function mutant of the Mp*FAD5* gene, which has enabled us to perform a comprehensive analysis into the role of HTA (16:3n3) and FAD5 in bryophytes. We show that C16 FAs and FAD5 are essential for proper chloroplast development in C16 plants. Additionally, we have further dissected the biosynthetic pathway of the bioactive hormone (defined as the ligand of COI1) in non-vascular plants, namely dn-OPDA. Our data indicate that, although the conversion of OPDA to dn-OPDA may exist in *Marchantia*, the majority of dn-OPDA is synthesised from HTA (16:3n3) *de novo*; i.e. almost exclusively via the 16:3, and not the 18:3, pathway. In the absence of HTA (16:3n3), dn-OPDA levels were extremely low in Mp*fad5* mutants. Surprisingly, however, typical dn-OPDA-mediated environmental stress responses could still be activated in the mutant. Results thus suggest the existence of other bioactive jasmonates, besides dn-OPDA, that activate the MpCOI1 receptor in *Marchantia*.

## RESULTS

### Identification of AtFAD5 ortholog in *Marchantia*

To determine if HTA (16:3n3) is a source of dn-OPDA in *Marchantia*, we first analysed the conservation of *FAD5*-related sequences in its genome. BlastP, sequence alignment and phylogenetic analysis revealed a single putative ortholog of At*FAD5* in *Marchantia*, corresponding to the gene *Mp3g10660*, which we subsequently named Mp*FAD5* (Sup. Fig. **S2**). To functionally characterize this gene, we generated loss-of-function mutants in the wild-type (WT; Tak-1) background using CRISPR-Cas9 Nickase technology (Fauser *et al*., 2014; Sugano *et al*., 2018). Two independent mutant lines, Mp*fad5#3* and Mp*fad5#8*, were selected for experimentation. Both lines lacked the first exon, including 80 base pairs upstream the starting ATG, and Mp*fad5#3* also lacked the first 30 base pairs of the second exon (Fig. **1a**). Phenotypic analysis of these mutants revealed paler green plants, with retarded growth compared to WT (Fig. **1b-f**). Analysis of photosynthetic pigments indicated a reduced level of total chlorophylls in Mp*fad5* mutants, which is consistent with the phenotype of At*fad5* mutants (Fig. **1c**). Moreover, measurement of F_v_/F_m_ (variable/maximum fluorescence of PSII; an indicator of the state of the photosystem), demonstrated a reduction in PSII photochemical efficiency in Mp*fad5* plants, indicative of a stressed or perturbed photosynthetic apparatus (Fig. **1d**). Mp*fad5* mutant plants also displayed a delay in antheridia formation after FR-mediated induction (Fig. **1d-f**). In some cases, plants did not generate sexual structures until one month after WT, and in others the antheridia initially appeared brownish, and finally aborted.

**Figure 1.**
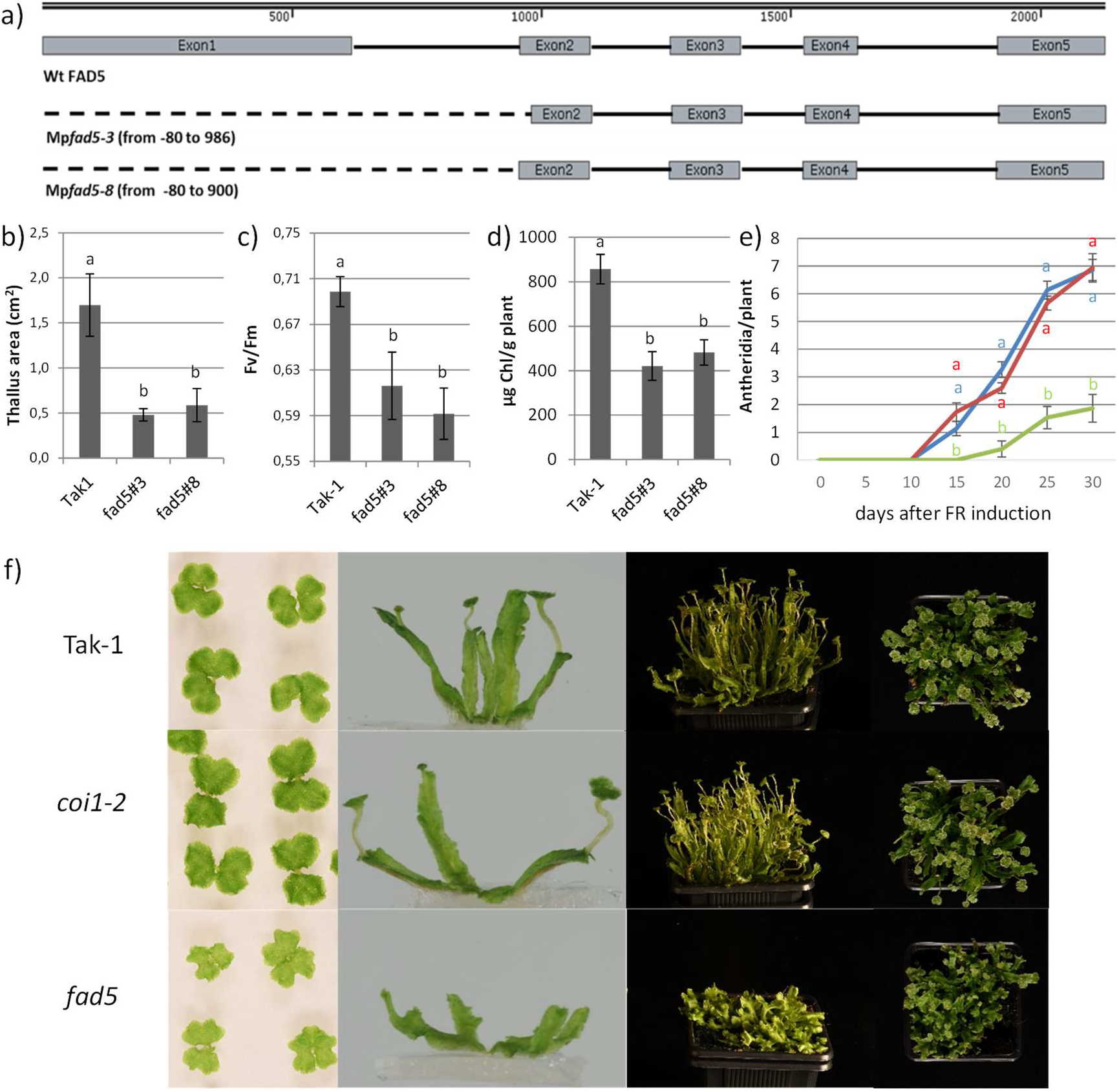
Mp*fad5* phenotype. **a)** Scheme of MpFAD5 gene. Grey blocks represent exons, dashed lines *fad5* lines deletions. **b)** Thallus area, **c)** Fv/Fm measured in dark adapted plants and **d)** total chlorophyll content after three weeks on culture. **e)** Timeline of number of antheridia per plant after FR induction (Tak-1: red line; Mp*coi1-2*: blue line; Mp*fad5*: green line). **f)** Tak-1, Mp*coi1-2* and Mp*fad5* plants growing for 2 weeks under constant light (first column) and after 15 days supplemented with FR (second column). Frontal and upper view of two months old plants after 30 days of FR supplementation (third and last column respectively). Data shown in **c)**, **d)**, **e)** and **f)** as mean ± S.D. (t-Tukey significant differences are shown as different letters).

### Mp*fad5* mutants display defects in the chloroplast structure

The lower PSII efficiency and the reduced growth rate of Mp*fad5* mutants suggested that they may have defective chloroplast function. Electron microscopy images of Mp*fad5* chloroplasts (compared to WT) revealed that their shape and internal organization were affected by the mutation. Although no differences in total number or size of chloroplasts were found (Sup. Table 1), the mutants had aberrant chloroplasts with a range of different shapes; from very narrow structures to those which were practically circular. This represents a marked difference to the classical oval shape of WT chloroplasts (Fig. **2**). The organization of thylakoid and grana also appeared to be compromised by the Mp*fad5* mutation. In the WT, clearly differentiated structures were observed forming parallel groups between themselves and the cell membrane. In Mp*fad5* cells, however, the thylakoid and grana conformation was disorganized and we found numerous cytoplasmic invaginations (Fig. **2**). These results indicate that FAD5 activity is essential for proper chloroplast development and function.

**Figure 2.**
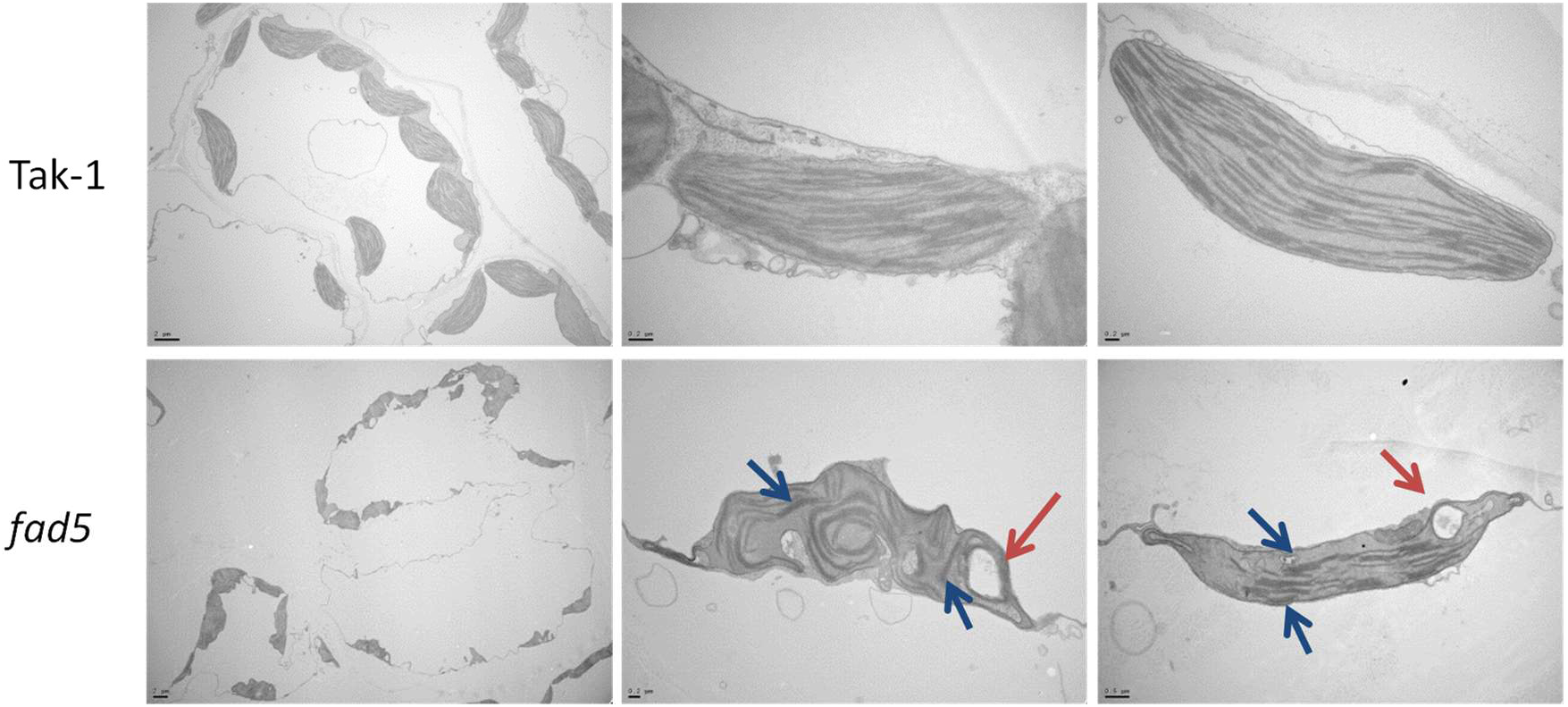
Chloroplast microscope analysis. Electronic microscopy images of Tak-1 and Mp*fad5* cells from the photosynthetic region of thallus (First column; Scale bars 2 μm). Detail of Tak-1 and Mp*fad5* single chloroplasts (Scale bars 0.2 μm, but last one of *fad5* 0.5 μm). Blue arrows indicate compressed grana. Red arrow indicates cytoplasm invagination.

### FAD5 is responsible for the synthesis of 16C PUFA and also affects the production of longer FAs

To establish whether the *FAD5* mutation affects the content of the main saturated and unsaturated fatty acids in *Marchantia*, we quantified prominent C16, C18 and C20 fatty acids in Mp*fad5* and WT plants by GC-MS. Mp*fad5* alleles were mainly impaired in the accumulation of unsaturated C16 FAs (Fig. **3a**). Neither HDA (16:2n6) nor HTA (16:3n3) were detected in Mp*fad5* alleles. A large reduction (70-80%, Fig. **3a**) of the peak corresponding to ∑16:1s was also observed. This peak is composed of two different 16:1 FAs that coelute; 7*Z*-hexadecenoic (HD; 16:1n9) and palmitoleic acid (16:1n7) (Kramer *et al*., 2002). Since FAD5 is a Δ^7^ desaturase (Heilmann *et al*.; 2004a) and the subproducts (HDA and HTA) of HD (16:1n9) were not detected, the reduction in this peak suggests the absence of HD, and the small observed amount of 16:1 should consist of palmitoleic acid (16:1n7). The contents of ALA (18:3n3) and eicosapentaenoic acid (EPA; 20:5n3) were also affected by the *FAD5* mutation (Fig. **3b**). Mp*fad5* plants displayed a reduction of around 40-50% for ALA and 25% for EPA compared with WT values. Altogether, these results indicate that MpFAD5 is responsible for the conversion of palmitic acid (16:0) into unsaturated C16:1n9 FA, which may contribute, at least partially, to the synthesis of longer FAs such as ALA and EPA.

**Figure 3.**
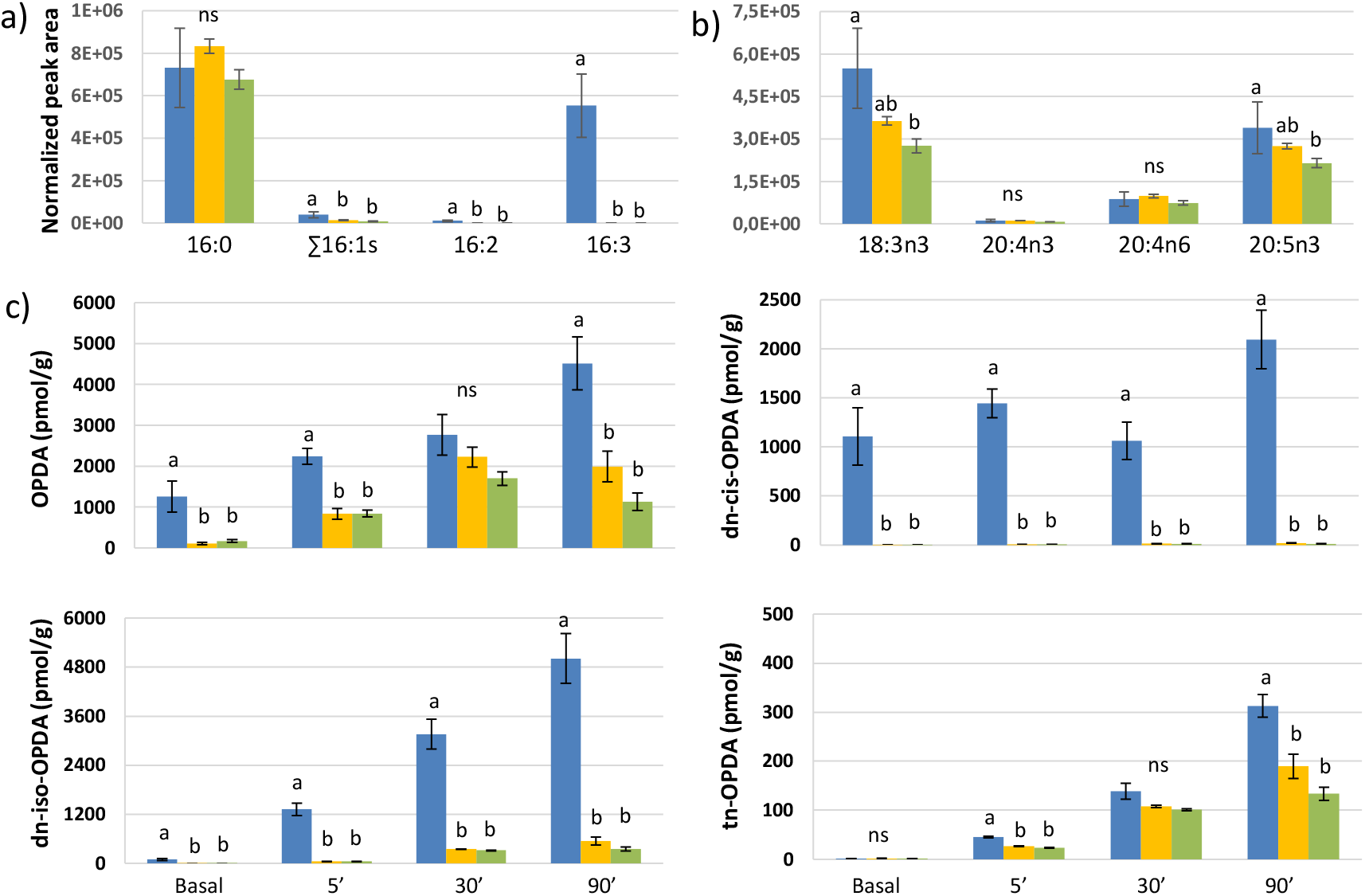
Fatty acids and jasmonates measurements. **a)** Content of C16 fatty acids (16:0 (palmitic acid), ∑ 16:1s [16:1Δ7 (7*Z*-hexadecenoic acid) + 16:1Δ9 (palmitoleic acid)], 16:2 (hexadecadoenoic acid, HAD) and 16:3 (hexadecatrienoic acid, HTA)) **b)** and some C18 and C20 fatty acids, 18:3n3 (α-linolenic acid; ALA), 20:4n3 (eicotetranoic acid; ETA), 20:4n6 (arachidonic acid; ARA) and 20:5n3 (eicosapentanoic acid; EPA) in Tak-1 (blue bars) and Mp*fad5* lines (yellow and green bars). **c)** Time-course accumulation of OPDA, dn-*cis*-OPDA, dn-*iso*-OPDA and tn-OPDA in Tak-1 (blue bars) and both Mp*fad5* mutant lines (yellow and green bars) in basal conditions and 5, 30 and 90 minutes after mechanical wounding. Data shown as mean ± S.D. (different letters mean significant differences t-Tukey test).

### Hexadecatrienoic acid is the main source of dn-ODPA in *Marchantia polymorpha*

To assess how changes in FA composition in Mp*fad5* affect the content of jasmonate-related oxylipins in *Marchantia*, we evaluated the accumulation pattern of these molecules after mechanical wounding in a time course experiment (Fig. **3c**). Significant differences were found between WT and Mp*fad5* lines for all measured jasmonates. Levels of both OPDA and tn-OPDA were seen to be moderately reduced, compared to WT. This finding was consistent with the reduction of ALA (18:3n3) in the Mp*fad5* mutant lines. The greatest differences, however, were observed in dn-*cis*- and dn-*iso*-OPDA contents. In Mp*fad5* mutants, dn-*iso*-OPDA levels reached only around 5% of that observed in WT plants, while dn-*cis*-OPDA was almost absent (less than 1%) in the mutant samples.

The strong reduction in dn-OPDA isomers correlate with the strong reduction in C16 FAs, whereas the smaller reduction in OPDA correlates with the reduction in ALA (18:3n3). Taken together, these results indicate that HTA (16:3n3) is the main source of both dn-OPDA isomers in *Marchantia*, contributing to ca. 95% of the total content. The contribution of OPDA to dn-OPDA production is, however, secondary and, at most 5%. In addition, the higher accumulation of dn-*iso*-ODPA (compared to dn-*cis*-) in the mutant, suggest that the kinetics of dn-*cis*- to dn-*iso*-OPDA conversion is faster than both the synthesis of dn-*cis*-ODPA from 16:3 and the β-oxidation of dn-*cis*-OPDA to produce tn-OPDA.

### dn-OPDA-mediated gene-expression is only partially compromised in Mpf*ad5*

Responses to mechanical wounding and insect feeding are among the main responses regulated by dn-OPDA isomers in *Marchantia* (Monte *et al*., 2018). To evaluate whether the reduction of dn-*cis/iso*-ODPA levels in Mp*fad5* alleles impairs responses to wounding, we obtained transcriptomic profiles of wounded *Marchantia* mutants and compared them with those of wounded WT plants. A total of 297 genes were differentially expressed (DE) by wounding in Mp*fad5* compared to WT plants (LogRatio > |1|, FDR<0.05). Clustering and gene ontology (GO) analyses comparing Mp*fad5* DE genes with previous data obtained in our laboratory (WT response to wounding, to OPDA treatment and DE genes in Mp*coi1* mutants in response to OPDA) revealed that most DE genes upregulated in Mp*fad5* were involved in chloroplast development and function, including responses to light (Clusters 4 and 5, Fig. **4a,b**). These results are consistent with the phenotypic defects in chloroplasts, chlorophylls content and PSII efficiency in Mp*fad5* alleles (Fig. **2**).

**Figure 4.**
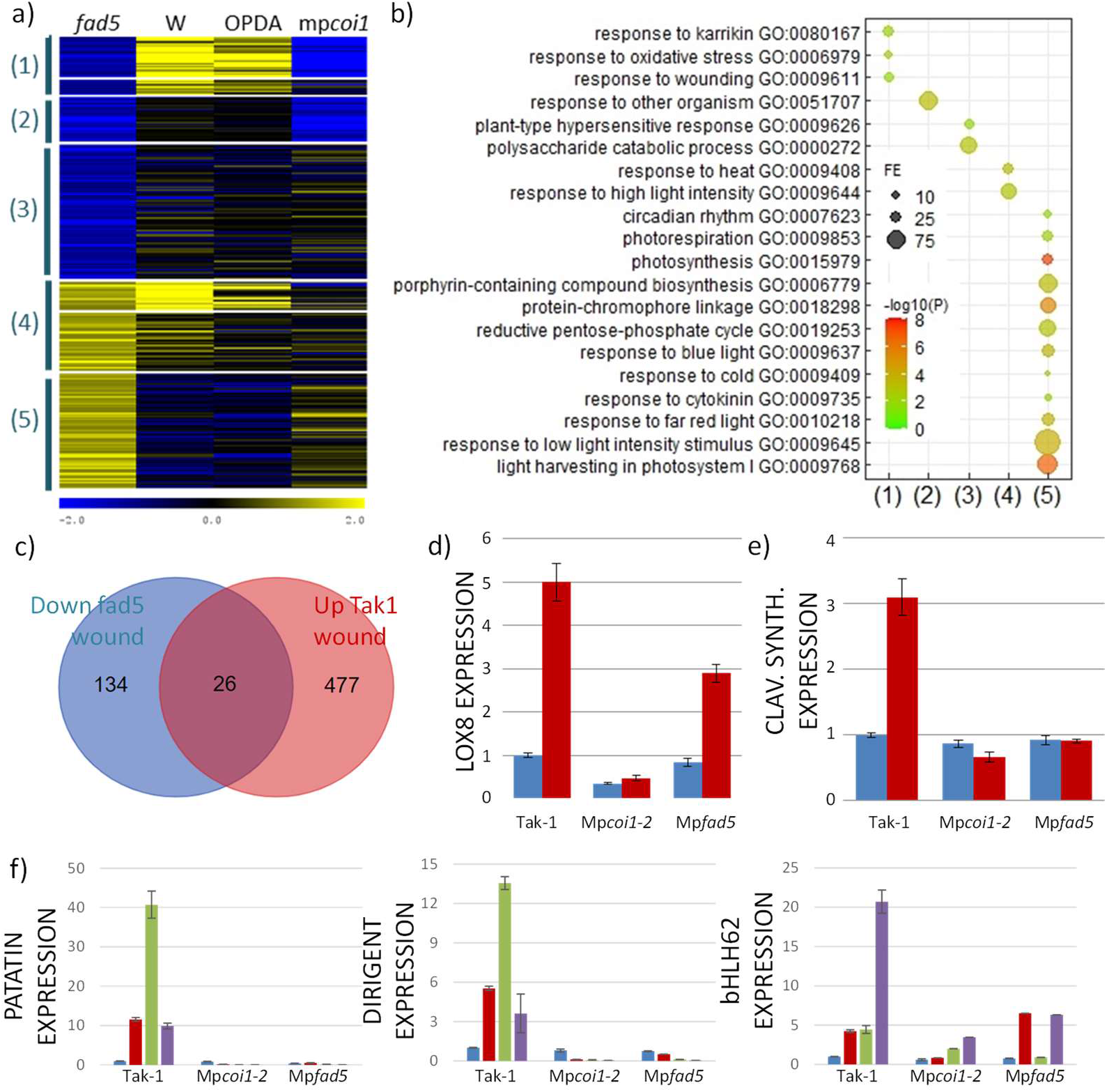
Gene expression analysis after different wounding stresses. **a)** K-means clustering of genes differentially expressed in response to wounding or OPDA in different genotypes. *fad5*: *fad5* wound vs Tak1 wound; W: Tak1 wound vs Tak1 mock; OPDA: Tak1 OPDA vs Tak1 mock; mp*coi1*, mp*coi1* OPDA vs Tak1 OPDA. Differentially expressed genes were selected from the *fad5* experiment (Log2[fad5/Tak1]>|1, FDR<0.05), and their expression in the rest of genotypes and treatments obtained from available data. Seven clusters were generated and some pairs with similar patterns were grouped for GO analyisis (numbers in brackets). **b)** Significant GO terms (biological process) associated to each of the groups in (b). FE, fold-enrichment of the term in the cluster. GO terms correspond to the most similar genes in Arabidopsis. **c)** Overlapping of genes down-regulated in *fad5* and up-regulated in Tak1 in response to wounding. Differentially expressed genes were selected as in (a). **d) e)** Q-PCR analysis of two COI dependent wound induced marker genes in basal conditions (blue bars) and after 90 minutes post wounding (red bars). **f)** Q-PCR analysis of three COI dependent wound induced marker genes in basal conditions (blue bars) and 1, 2 or 4 days after wounding (red, green and purple bars respectively).

Further to this, our analyses identified a group of the downregulated genes in Mp*fad5* to be genes regulated by wounding, OPDA or MpCOI1 (Clusters 1 and 2; Fig. **4a**). The top section of cluster 1 represents genes regulated by wounding, OPDA and MpCOI1, whereas the bottom part represents genes regulated by wounding and OPDA, but independent of MpCOI1. Cluster 2 represents genes regulated by MpCOI1 but not by wounding or OPDA. GO and qPCR analyses confirmed our data. The GO term “response to wounding” was enriched in cluster 1, whereas “response to other organisms” was enriched in cluster 2 (Fig 3B; lanes 1 and 2). The qPCR analysis of early and late dn-OPDA-responsive marker genes (Mp*PATATIN*, Mp*DIRIGENT*, Mp*bHLH62*, Mp*LOX8* and Mp3g11070 (*Clavaminate synthase-like*)) showed marked differences in wound-mediated induction between WT and Mp*fad5*, consistent with the wound/OPDA regulation of genes in both clusters 1 and 2, and further supported that genes in cluster 2 may represent late wound-responsive genes (Fig. **4c**). These results support the importance of dn-OPDA and MpFAD5 in the MpCOI1-dependent response to wounding.

Surprisingly, however, when considering the huge number of genes regulated by wounding in WT plants (Fig. **4c**), the relative contribution of MpFAD5 to wound-related responses was much lower than we expected. Indeed, among the 503 genes induced by wounding in WT plants, only 26 were DE in Mp*fad5* plants. This result could be explained by the small quantity of dn-*iso*-OPDA accumulating in Mp*fad5* (Fig. **3c**). However, given that around 95% of normal dn-*iso*-OPDA levels are lost in this mutant, this data rather suggests that dn-OPDA is not the only molecule activating the wound response in *Marchantia*.

### Mp*fad5* plants have a WT-like response to herbivory

To further study the consequence of reduced dn-OPDA isomers and the role of MpFAD5 in defence, Mp*fad5* mutants were subjected to herbivory using WT and Mp*coi1-2* plants as control. First instar larvae of the generalist herbivore *Spodoptera exigua* were released on 6-week-old Marchantia plants and their performance (larval weight) was measured after seven days. In agreement with previous results (Monte *et al*., 2018), larvae fed on Mp*coi1-2* were significantly larger than those collected from WT plants (Fig. **5a**). However, and despite the extremely low levels of dn-OPDA, those fed on Mp*fad5* plants were not significantly different to those fed on WT plants, indicating that defences against herbivores are not impaired in Mp*fad5*.

**Figure 5.**
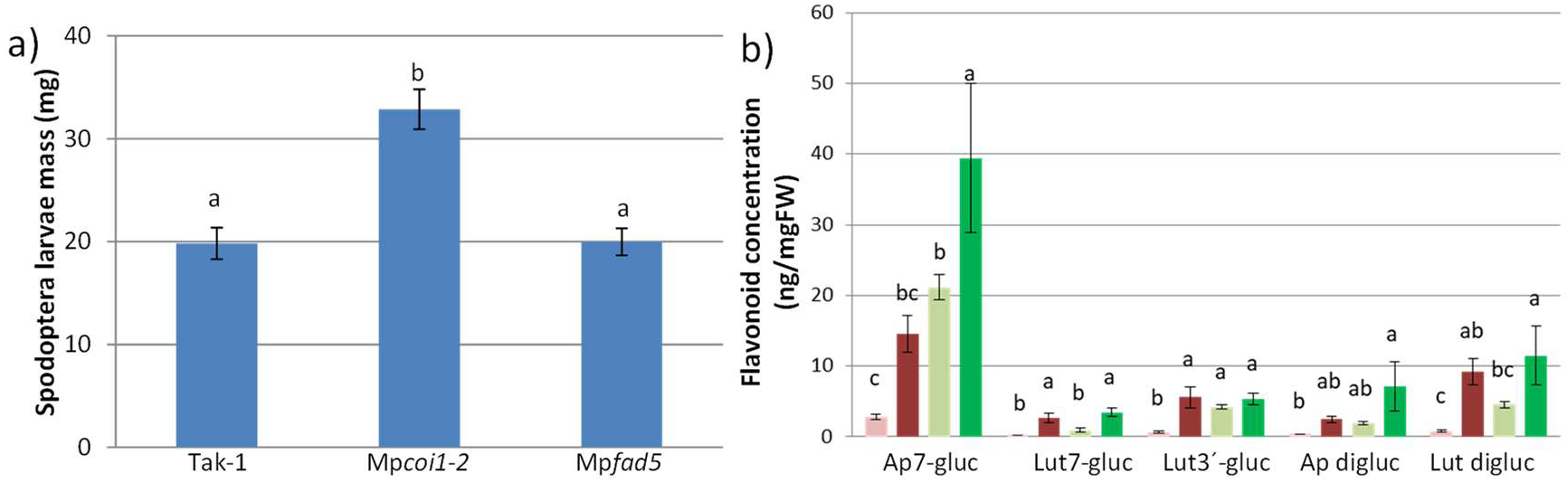
Larval infection. **a)** *Spodoptera exigua* larval weight after 9 days feeding on *M. polymorpha* thalli of Tak-1, Mp*coi1-2* and Mp*fad5*. **b)** Content of main flavonoids found in *M. polymorpha* (Tak-1: reddish bars, and Mp*fad5*: green bars) in mock (paler bars) and after herbivory stress (darker bars). (Ap7-gluc, apigenin 7-*O*-glucuronide; Lut7-gluc, luteolin 7-*O*-glucuronide; Lut3’-gluc, luteolin 3’-*O*-glucuronide; Ap digluc, apigenin 7,4’-di-*O*-glucuronide; Lut digluc, luteolin 7,3’-di-*O*-glucuronide). Data shown as mean ± S.E in **(a)** and as mean ± S.D in **(b)** (different letters mean significant differences t-Tukey test).

Since light stress induces flavonoid accumulation in *Marchantia* (Albert *et al*., 2018; Soriano *et al*., 2019) and flavonoids are known to be involved in defence against herbivores (Morimoto *et al*., 2003; Qi *et al*., 2008), we analysed the flavonoid contents in Mp*fad5* after herbivory employing an experimental set-up similar to that in Fig. **5a** (seven days after treatment). *Marchantia* has previously been shown to accumulate mainly apigenin and luteolin derived flavones (Markham & Porter, 1974; Soriano *et al*., 2019; 2021). As shown in Fig. **5b**, higher levels than WT for all measured flavonoids in basal conditions. Similarly, after herbivory, Mp*fad5* showed higher values of apigenin derivatives (apigenin 7-*O*-glucuronide and apigenin 7,4’-di-*O*-glucuronide) and also for one luteolin (luteolin 7’-*O*-glucuronide), in comparison to WT plants.

These results show that despite severe reduction in dn-OPDA levels in the absence of MpFAD5, the plant defences are unaffected. Given that only minor perturbations of the wound-responses were seen, taken together, our data is strongly indicative of an alternate signalling molecule that may act redundantly with dn-OPDA in the activation of jasmonate responses.

## DISCUSSION

Despite the importance of dn-OPDA as the bioactive jasmonate, i.e. the COI1 ligand in *Marchantia*, details of its biosynthesis in bryophytes are still unclear. Previous work in *Arabidopsis* suggested two possible routes for dn-OPDA production: the hexadecanoid pathway, producing dn-OPDA directly from HTA (16:3n-3), and the octadecanoid pathway, producing the molecule indirectly from ALA (18:3n3, via an OPR3-independent conversion of OPDA to dn-OPDA (Weber *et al*., 1997; Chini *et al*., 2018). In *Marchantia*, exogenous treatment with labelled ALA (18:3n3) and HTA (16:3n3) confirmed that both FAs are potential dn-OPDA precursors (Monte *et al*., 2018), though their relevance *in vivo* was not addressed. Here, by analysing Mp*fad5* mutants, we show that although some conversion of OPDA into dn-OPDA is possible, particularly after exogenous treatment, the main source of dn-OPDA in *Marchantia* is the hexadecanoid pathway [i.e. HTA (16:3n-3)], and the contribution of the octadecanoid pathway is minimal. Moreover, we show the functional conservation of FAD5 in species separated by more than 450 million years of independent evolution, suggesting that this function evolved in the common ancestor of all land plants.

Similar to At*fad5* (Heilmann *et al*., 2004a), Mp*fad5* mutants lack Δ^7^-desaturase function and, therefore, are not able to synthesize HD (16:1n9), HAD (16:2n6) and HTA (16:3n3). In contrast, 18:1 and 18:2 levels are unaffected and 18:3 levels remain high in Mp*fad5*.The consequence of blocking the hexadecanoid pathway in Mp*fad5* is a reduction of around 95% of the dn-OPDA content and virtually 100% of the dn-cis-OPDA isomer. Thus, the octadecanoid pathway, and the conversion of OPDA into dn-OPDA, is clearly a secondary and minor source for the synthesis of this compound, accounting for, at most, 5% of the total dn-OPDA content *in vivo*.

Blockage of the hexadecanoid pathway in Mp*fad5* is characterized by the absence of 16:2 and 16:3 in this mutant. However, MpFAD5 catalyses the unsaturation of 16:0 to 16:1 and the peak corresponding to 16:1 does not fully disappear in the mutants. This peak is composed of two different 16:1 FAs that coelute; 7*Z*-hexadecenoic (HD; 16:1n9) and palmitoleic acid (16:1n7) (Kramer *et al*., 2002). Since FAD5 is a Δ^7^-desaturase (Heilmann *et al*.; 2004a) and the subproducts (HDA and HTA) of HD (16:1n9) were not detected, the presence of this peak can be attributed to palmitoleic acid, with the reduction in the peak size being caused by the absence of HD. *Marchantia* has 3 different orthologs of *Arabidopsis* FAB-family member FAB2, one of which displays high similarity to a Δ^9^-desaturase involved in palmitic acid synthesis in the diatom *Phaeodactylum tricornutum* (Liu *et al*., 2020). The remaining content of palmitoleic acid (16:1) could therefore be explained by the actions of this MpFAB2.

Despite the functional conservation between AtFAD5 and MpFAD5, differences in FA profile between Mp*fad5* and WT plants suggest that FAD5 function in *Marchantia* may be more extensive than that in *Arabidopsis*. Whilst 16C FAs in At*fad5* mutants are clearly reduced, 18C FAs were found to be unaffected (Heilmann *et al*., 2004a). In contrast, our data demonstrates that in Mp*fad5*, 18:3n3 levels are lower than in WT, as are levels of the longer chain FA, 20:5n3 (which is not present in *Arabidopsis*). The wider-reaching effects of MpFAD5 on longer chain FAs may be explained by divergences in FAD5 function between *Arabidopsis* and *Marchantia*; changes that have likely resulted from differences in the relative contribution of other proteins involved in FA biosynthesis. Indeed, one of the main differences between the genomes of these two species is the level of gene redundancy, which is much greater in *Arabidopsis* compared with *Marchantia*. In this instance, this is hugely relevant since the ADS (*Arabidopsis* Desaturase) family, which is numerous in *Arabidopsis* (including 8 ADSs besides AtFAD5), is greatly diminished in *Marchantia*, which has only one ADS member: MpFAD5 (Sup. Fig. **S2**). Previous studies on AtADSs have shown that their regiospecificity varies according to their intracellular location (i.e. they act as Δ^7^ in the plastids and as Δ^9^ in the cytoplasm) and that consequently they can desaturate not only palmitic acid but also stearic acid (Heilmann *et al*., 2004b). Thus, being the only member of the ADS family in *Marchantia*, MpFAD5 may integrate several of the functions distributed across distinct genes in *Arabidopsis* and consequently can participate in the desaturation of both types of FAs (C16 and C18). As such, it appears that a significant part of the evolution of the ADS family in *Arabidopsis* has involved functional specialization.

The analysis of Mp*fad5* mutants also uncovered that the deficit in unsaturated 16C FAs significantly affects the shape and function of chloroplasts. In addition to changes in chloroplast morphology, other markers of the chloroplasts being in poor conditions included compressed grana and the presence of unusual invaginations (Woodson, 2019; Alamdari *et al*., 2021; Lemke *et al*., 2021). This defect in the structure of chloroplast membranes results in a poor performance of the photosynthetic apparatus. F_v_/F_m_ values of Mp*fad5* mutant plants indicate that these plants were stressed (photoinhibited) in basal growth conditions. Previous studies in *Marchantia* have shown that F_v_/F_m_ values depend on light conditions, with lower values (photoinhibited) at high radiation (Soriano *et al*., 2019). Gene-expression analysis confirmed a constitutive response to light stress, a likely consequence of the defects in chloroplasts development.

Developmental defects in the photosynthetic apparatus resulted in a lower growth rate, a pale green phenotype and other anomalous light responses such as the delay in archegonia formation (regulated by Far-Red light; Inoue *et al*., 2019). Notably, the pale green phenotype is also characteristic of the At*fad5* mutant in *Arabidopsis* and of tomato *spr2* mutant, which cannot synthesize HTA (Heilmann *et al*., 2004a; Li *et al*., 2003). HTA (16:3n3) is one of the main components of monogalactosyl-diacylglycerols (MGDGs) in the so-called 16:3 plants (those with a predominance of the prokaryotic route, such as *Arabidopsis* and *Marchantia*; Reszczyńska & Hanaka, 2020). These, together with digalactosyl-diacylglicerols (DGDGs), containing mainly 18:3 FAs, are the predominant lipids found in thylakoid membranes. Although chloroplast development has not been analysed in At*fad5* or Sl*spr2*, the phenotypic coincidence in these evolutionary distant species strongly suggests that HTA (16:3n3) is essential for chloroplasts membrane integrity and organelle development in 16:3 plants.

Besides a strong activation of chloroplasts and light stress related genes in Mp*fad5*, our transcriptomic analyses revealed a defect in the induction of a subset of dn-OPDA regulated genes, which are activated by MpCOI1 and the jasmonate pathway in response to wounding. Whilst these data support a role for MpFAD5 and HTA (16:3n3) in dn-OPDA biosynthesis and signalling, it was surprising to observe that the deficit in the transcriptional response to mechanical wounding was not much greater. Given the extent to which dn-OPDA levels are reduced in Mp*fad5* mutant plants, it is somewhat remarkable that the majority of the wound-responsive genes were unaffected in this line. Although it is possible that wound-related gene activation is a result of the small quantity of dn-*iso*-OPDA accumulating in Mp*fad5*, similar levels (ca 5%) of the vascular plant hormone, JA-Ile, were sufficient to activate a much stronger jasmonate-response in the *Arabidopsis* mutant *opr3* (Chini *et al*., 2018). Similarly, At*opr3* plants showed an increased susceptibility to herbivorous insects compared to WT, whereas Mp*fad5* displayed WT levels of resistance. One explanation for this is the increased level of flavonoids, such as apigenin and luteolin derivatives, that are constitutively found in Mp*fad5* mutants. These flavonoids have been shown to have antifeedant activity (Golawska & Lukasik, 2012; El Shafeiy & Abdelaziz, 2020) and thus may have contributed to the protection of the mutant plants. Furthermore, it has been found that ω3 VLPUFA (Very Long Chain PUFA) favours larvae development (Twining *et al*., 2016), and therefore, the reduced levels in Mp*fad5* may also contribute to the mutant plants’ success against herbivorous insects.

Nonetheless, the near-normal level of jasmonate-responsive gene expression in Mp*fad5* mutants does suggest that the resistance to herbivory in these plants cannot be explained by flavonoid content and reduced ω3 VLPUFA levels alone. Moreover, in addition to many wound-responsive genes being unaffected in Mp*fad5* plants, a subset of dn-OPDA regulated heat response genes were found to be upregulated. HS genes are specifically regulated by dn-*cis*-OPDA (and not by dn-*iso*-OPDA, which lacks electrophilic properties; Monte *et al*., 2020), lending weight to the argument that the largely unaffected jasmonate-related responses in Mp*fad5* are not solely a result of the small amount of dn-*iso*-OPDA which accumulates.

Given this, in conjugation with the different effects that similar reduction in hormone levels have in Mp*fad5* and At*opr3*, it is tempting to speculate that dn-OPDA is not the only jasmonate activating wound responses in *Marchantia*. Our data strongly suggest the existence of an as-of-yet undiscovered molecule(s) which may act synergistically with dn-OPDA to activate defence responses against both mechanical wounding and herbivory. *Marchantia* membranes contain several VLPUFAs (Lu *et al*., 2019), which are not common in vascular plants, and may act as a source for alternate signalling molecules in the jasmonate pathway. Since the study of jasmonates in plants besides *Arabidopsis* has, until recently, been largely absent in the scientific literature, understanding the contribution of VLPUFAs to jasmonate signalling has been greatly overlooked. Our findings indicate that new branches of the jasmonate pathway and clues to its evolution are hidden within different plant species, and presents an exciting future challenge for the field.

## MATERIAL AND METHODS

### Plant material

*Marchantia polymorpha* accession Takaragaike-1 (Tak-1) was used as WT. In this genetic background, we used CRISPR–Cas9_D10A_ nickase-mediated mutagenesis with Mp*FAD5* as the target. The sequence of this gene was obtained from www.marchantia.info. Four different gRNAs were designed flanking the first exon of the gene. Then they were cloned firstly into pBC-GE12, pBC-GE23, pBC-GE34 and pMPGE_EN04 vectors, to subsequently be transferred in tandem by LR reaction to the pMpGE018 binary vector carrying the CRISPR–Cas9_D10A_ nickase (Ran *et al*., 2013; Shen *et al*., 2014). Using regenerating thalli transformation (Kubota *et al*., 2013), Tak-1 plants were transformed and thalli selected by chlorosulfuron resistance. Then, gDNA of transformants were extracted and sequenced using gRNAs flanking primers (Sugano & Nishihama, 2018). In addition to the mutants generated in this study, Mp*coi1-2* (Monte *et al*., 2018) was used in some experiments as a control for dn-OPDA insensitivity.

### Culture conditions

Plants were routinely grown at 21°C, under continuous white light (50-60 mmol m^-2^ s^-1^) on Petri plates containing 1% agar half-strength Gamborg’s B5 medium. For fatty acids quantification, wounding assays (both hormone measurements and gene expression), chlorophyll extraction and chlorophyll fluorescence measurements plants were grown for three weeks.

For the antheridiophore induction experiment, gemmae of Tak-1, Mp*fad5* and Mp*coi1-2* were placed in Gamborg’s B5 plates and after ten days transferred to soil. Three weeks after transplant, promotion of antheridiophores was induced by supplementation of white light with far red (Chiyoda *et al*., 2008; Kubota *et al*., 2014).

### Pigment contents and photosynthetic variables

Maximum (F_m_) and minimum (F_0_) chlorophyll fluorescence values were measured after 21 days of culture, using a portable pulse amplitude modulation fluorometer (MINI-PAM, Walz, Effeltrich, Germany). Then, the maximum quantum yield of PSII (F_v_/F_m_) was determined, where F_v_ = F_m_ − F_0_.

Photosynthetic pigments were extracted with 95% ethanol after freezing shoot apices in liquid N_2_ and grounding them in a TissueLyser (Qiagen, Hilden, Germany). Chlorophylls a and b were quantified by spectrophotometry (Sumanta *et al*., 2014).

### Microscopy analysis of chloroplast

*Marchantia* thalli were cut in small pieces (1×2 mm) and fixed immediately in buffered phosphate 0,1 M paraformaldehyde 4% (Electron Microscopy Sciences), glutaraldehyde 2,5% (TAAB Laboratories) for 3h, room temperature and for 48h, 4°C. Samples were washed with phosphate sucrose buffer 0,1M, post-fixed (1h, 4°C) with 1 % osmium tetroxide (TAAB Laboratories) in potassium ferricyanide 0,8% (Sigma) and incubated with 2% aqueous uranyl acetate (Electron Microscopy Sciences) for 1h at 4°C. After washing with distilled water, samples were dehydrated with increasing concentrations of acetone (anhydrous, VWR) and embedded in epoxy resin TAAB 812 (TAAB Laboratories). Polymerization was carried out (in epoxy resin 100%) for 2 days, 60°C. Resin blocks were detached and ultrathin 70 nm-thick sections were obtained with the Ultracut UC6 ultramicrotome (Leica Microsystems), transferred to 200 mesh formvar carbon coated nickel grids (Gilder) and stained with 2% aqueous uranyl acetate (30 min) and lead citrate 0,2% (1 min) at room temperature. Sections were visualised on a JEOL JEM 1011 electron microscope operating at 100 kV. Micrographs were taken with a Gatan Erlangshen ES1000W digital camera at various magnifications.

### Phytohormone and fatty acids measurements

3 weeks old thalli of Tak-1, Mp*fad5-3, Mpfad5-8* were mechanically wounded with tweezers. After 5, 30 and 90 minutes, wounded thalli were harvested and immediately frozen and ground in liquid nitrogen. Four replicates of each genotype and time point were collected. Endogenous *cis*-ODPA, dn-*cis*-OPDA, dn-*iso*-OPDA, tn-*cis*-OPDA, tn-*iso*-OPDA, were analysed using high performance liquid chromatography electrospray-high-resolution accurate mass spectrometry (HPLC–ESI–HRMS). For hormone extraction and purification, ±100 mg of frozen powder was homogenized with 1 ml precooled (−20 °C) methanol:water:HCOOH (90:9:1, v/v/v with 2.5 mM Na-diethyldithiocarbamate) and 10 μl of a stock solution of 1,000 ng ml^-1^ of the deuterium-labelled internal standards [^2^H_5_]dnOPDA and [^2^H_5_]OPDA in methanol. Samples were extracted by shaking in a Multi Reax shaker (Heidolph Instruments) (60 min at 2,000 r.p.m. at room temperature). After extraction, solids were separated by centrifugation (10 min at 20,000g at room temperature) in a Sigma 4-16K Centrifuge (Sigma Laborzentrifugen), and re-extracted with an additional 0.5 ml extraction mixture, followed by shaking (20 min) and centrifugation. 1 ml of the pooled supernatants was separated and evaporated at 40 °C in a RapidVap Evaporator (Labconco Co). The residue was redissolved in 250 μl methanol/0.133% acetic acid (40:60, v/v) and centrifuged (10 min, 20,000 RCF, room temperature) before injection into the HPLC–ESI–HRMS system. Hormones were quantified using a Dionex Ultimate 3000 UHPLC device coupled to a Q Exactive Focus Mass Spectrometer (Thermo Fisher Scientific) equipped with an HESI(II) source, a quadrupole mass filter, a C-trap, a HCD collision cell and an Orbitrap mass analyser, using a reverse-phase column (Synergi 4 mm Hydro-RP 80A, 150 × 2 mm; Phenomenex). A linear gradient of methanol (A), water (B) and 2% acetic acid in water (C) was used: 38% A for 3 min, 38% to 96% A in 12 min, 96% A for 2 min, and 96% to 38% A in 1 min, followed by stabilization for 4 min. The percentage of C remained constant at 4%. Flow rate was 0.30 ml min-1, injection volume was 40 μl, and column and sample temperatures were 35 and 15 °C, respectively. Standards of *cis*-12-oxo-phytodienoic acid (OPDA) and [^2^H_5_]OPDA were purchased from OlChemim Ltd (Olomouc, Czech Republic), and standards of dinor-12-oxophytodienoic acid (dn-OPDA) and [^2^H_5_]dnOPDA were purchased from Cayman Chemical Company (Ann Arbor, MI, USA). dn-*iso*-OPDA and tn-*iso*-OPDA were synthesized as previously described (Monte *et al*., 2018; Chini *et al*., 2018).

For FA measurements, samples of 3 week old plants were collected, freeze dried, and extracted (ca. 10 mg, exact mass was recorded) and analized as previously described (Ichihara & Fukubayasi, 2010). Nonadecanoic acid was used as the internal standard. GC-MS analysis was performed in an Agilent QTOF-A 7890B GC System coupled to a 7250 Accurate Mass Q-TOF and a 7693A Autosampler. The temperature program was as follow: The injection volume was 0.5 μL in an Agilent GC column DB5-MS (30 m length, 0.25 mm i.d, and 0.25 μm film of 95% dimethyl/5% diphenylpolysiloxane) with a precolumn (10 m J&W integrated with Agilent 122-5532G) was used for compound separation. Carrier gas flow rate (He) was set at 1.9852 mL/min and the split/splitless injector and transfer line temperatures at 250 °C and 280 °C, respectively and the split ratio was 1:5. The initial column oven temperature was set at 80 °C (held for 1 minute), rose to 200 °C at 15 °C/min, to 280°C at 10°C/min, to 325°C at 25°C/min and hold this temperature 7 min before cooling down for the next injection. The total run time was 25.8 min. MS detection was performed with electron impact ionization (EI) at −70 eV of energy and 250 °C in filament source.

### Gene expression analysis

A custom Marchantia microarray was used for Tak-1 and Mp*fad5* mutant lines (see Monte *et al*., 2018). Three weeks old thalli were wounded with tweezers, collected after 90 minutes and their RNA extracted with FavorPrep Plant Total RNA Mini Kit. Contaminating DNA was removed by DNase on-column digestion and sample quality was tested in a Bioanalyzer 2100 (Agilent Technologies). Four independent biological replicates of wounded Tak-1 and Mp*fad5* plants were processed.Hybridizations, slide scanning and bioinformatic processing were performed as in Monte *et al*.(2018). Differentially expressed genes were selected by log-ratio values above 1 and below −1 and with an expected false discovery rate, FDR (Limma) < 5%. Clustering of genes was performed using K-Means with Euclidean distance (Soukas *et al*., 2000) in Multi Experiment Viewer (http://mev.tm4.org/).

### Herbivory assays

Plates with 7 plants of Tak-1, *coi1-2*, and *fad5* were grown as described above for 2 weeks. After this period, they were transferred to soil for other three weeks. Then, 10 first instar larvae of *S. exigua (Entocare, The Netherlands)* were released in each pot with 6 weeks old plants. 5 pots were used for each genotype. To keep the insects in the pot, they were covered with perforated plastic boxes. After one week of feeding, larvae were collected and weighed on a precision balance.

### Individual flavonoids measurements

Concentration of main flavonoids found in *Marchantia polymorpha* were measured by ultra-performance liquid chromatography (UPLC) using a Waters Acquity UPLC system (Waters Corporation, Milford, MA, USA), following Soriano *et al*. (2019). The UPLC system was coupled to a micrOTOF II high-resolution mass spectrometer (Bruker Daltonics, Bremen, Germany) equipped with an Apollo II ESI/APCI multimode source and controlled by the Bruker Daltonics Data Analysis software. Standars of apigenin and luteolin were used to quantify the five compounds (Sigma-Aldrich, St. Louis, MO, USA).

## ACKNOWLEDGEMENTS

We thank Grupo de Ecofisiología Vegetal of Universidad de La Rioja (Ecophys) for access to UPLC analysis of flavonoids. We also thank the Electron Microscopy Facility (Centro Nacional de Biotecnologia, Universidad Autonoma, Madrid) for preparing samples (Epon embedding), obtaining the ultrathin sections and TEM visulaization. GS was supported by a postdoctoral grant funded by Universidad de La Rioja. SK was supported by an EMBO Long-Term fellowship (ALTF 47-2017) and a Juan de la Cierva fellowship from the Spanish Ministery for Science and education (IJC2018-035580-I) GH. J-A. was supported by the Deutsche Forschungsgemeinschaft (Individual Research Grant JI 241/2-1). This work was funded by the Spanish Ministry for Science and Innovation grant PID2019-107012RB-100 (MICINN/FEDER) to R. Solano. JMFZ’s lab was funded by grant BIO2017-86651-P (MINECO/FEDER).

## AUTHOR CONTRIBUTION

G.S., S.K., G.J-A and R.S. conceptualized and designed the research. G.S., S.K., G.J-A conducted all experiments and analyzed data. A.M.Z., J.M. G-M, F. R-S. V. and C.B. made all metabolite measurements. J.M. F-Z. made transcriptomic analyses. G.S., S.K., G.J-A and R.S wrote the manuscript with input from all co-authors. All authors reviewed and approved the manuscript.

## DATA AVAILABILITY

All data will be available upon request to R.S.

## SUPPLEMENTAL INFORMATION

**Supplemental Figure S1:**
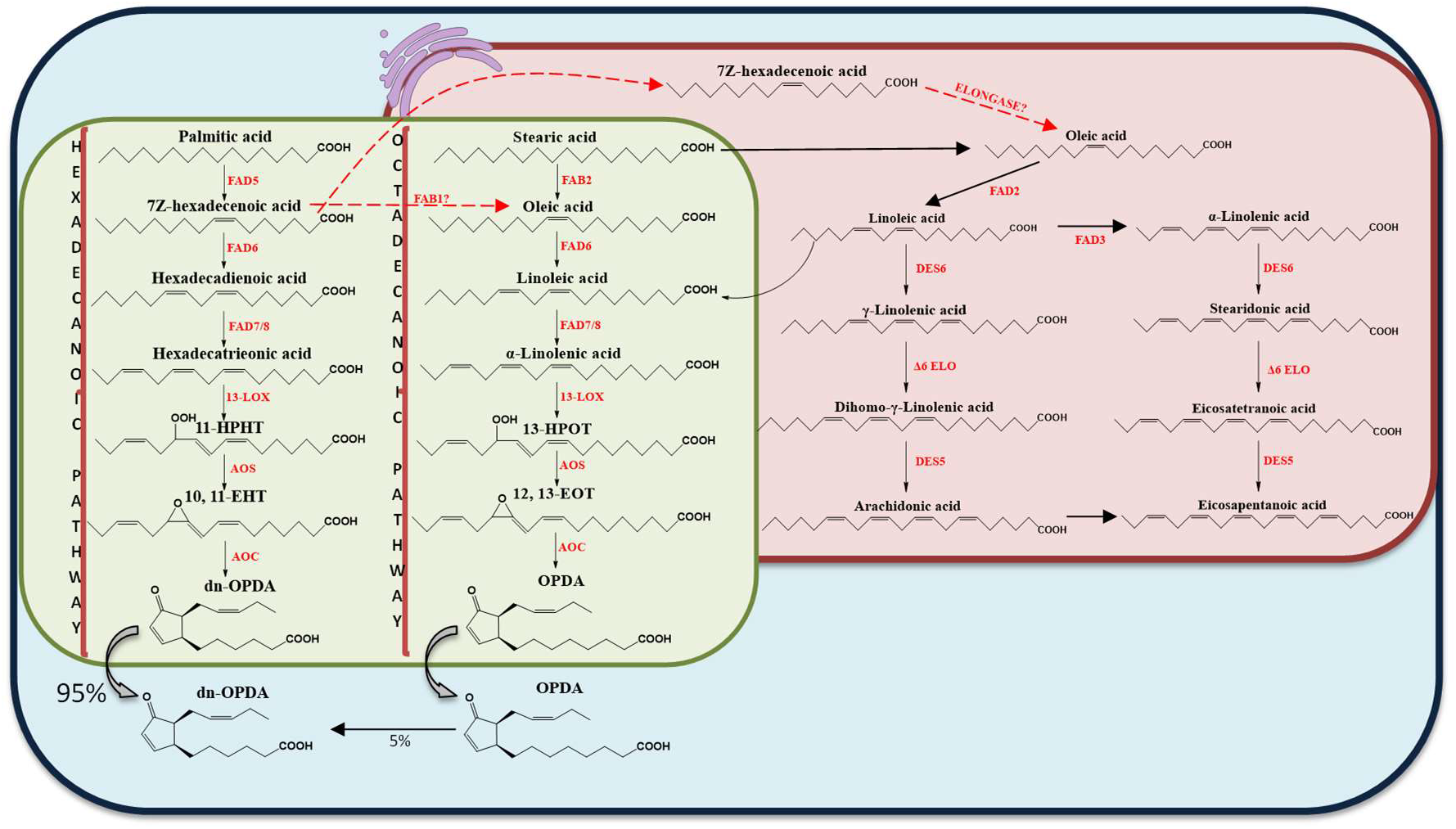
Scheme of ODPA and dn-OPDA biosynthesis. OPDA and dn-OPDA biosynthesis occurs in the chloroplast (green) where α-linolenic acid (18:3n3), octadecanoid pathway, or hexadecatrienoic acid (16:3n3), hexadecanoid pathway, are released from chloroplast membranes. Then, they are subjected to the consecutive actions of lipoxygenase, allene oxide synthase (AOS) and allene oxide cyclase (AOC) enzymes, to produce ODPA and dn-OPDA.

**Supplemental Figure S2.**
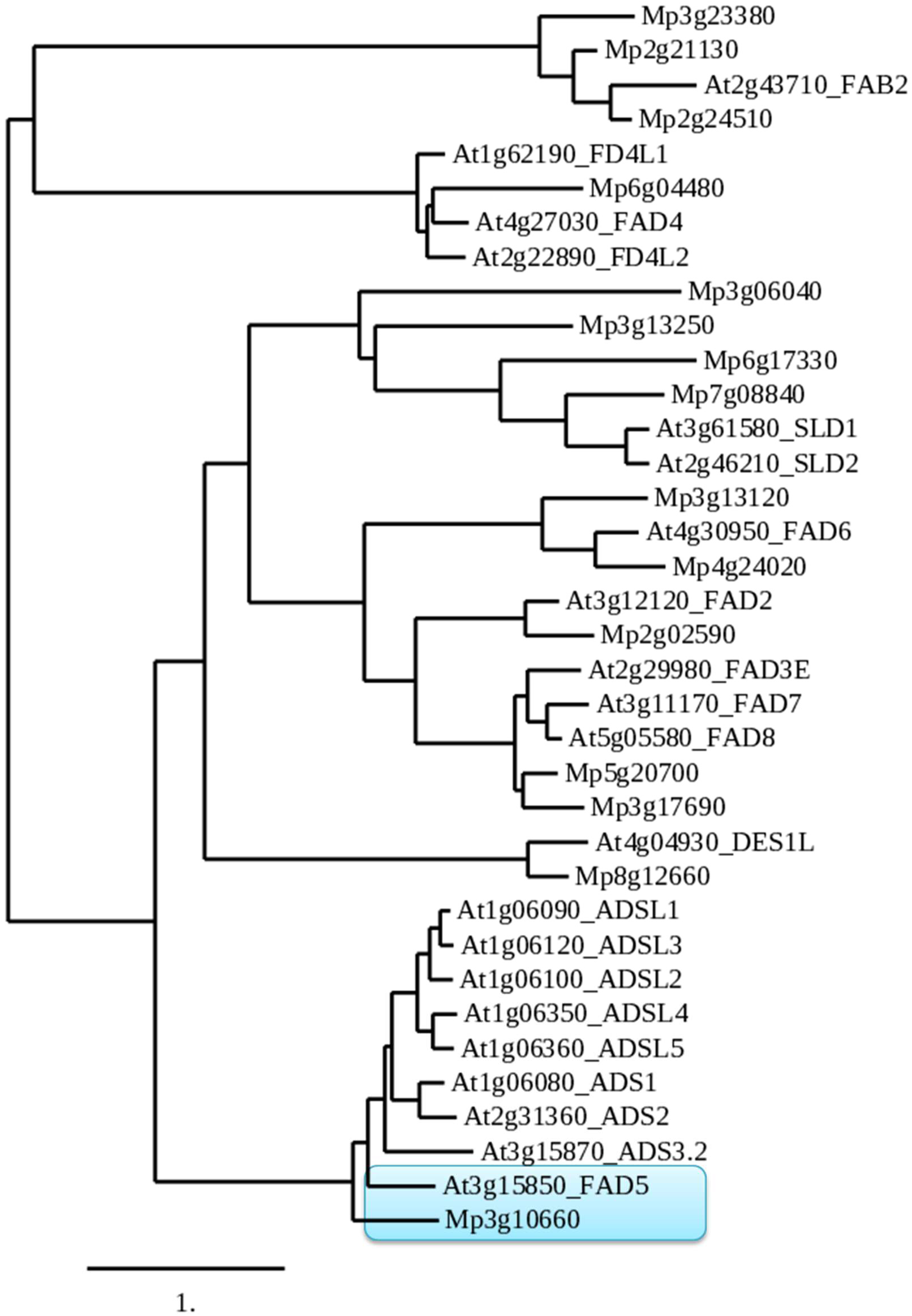
Phylogenetic tree of *Arabidopsis thaliana* and *Marchantia polymorpha* desaturases proteins. By using http://www.phylogeny.fr/, amino acid sequences were aligned by MUSCLE and tree was generated using the neighbour-joining method.

**Supplemental Table 1.** Size measurements (Mean ± S.D.) of Tak-1 and Mp*fad5* chloroplasts (n=39) made with ImageJ software.

|  | Length (μm) | Width (μm) | Area (μm<sup>2</sup>) |
| --- | --- | --- | --- |
| Tak-1 | 5,34 ± 0,31 | 1,82 ± 0,13 | 8,40 ± 0,74 |
| Mpfad5 | 5,64 ± 0,33 | 1,74 ± 0,16 | 7,33 ± 0,69 |

